# Reduced fibrous capsule elastic fibers from biologic ECM-enveloped CIEDs in minipigs, supported with a novel compression mechanics model

**DOI:** 10.1101/2022.08.01.502321

**Authors:** Roche C. de Guzman, Allison S. Meer, Aidan A. Mathews, Atara R. Israel, Michael T. Moses, Clarence M. Sams, Daniel B. Deegan

**Author notes:** Corresponding author: Roche C. de Guzman, 133 Hofstra University, 203 Weed Hall, Hempstead, NY 11549, USA.

## Abstract

**BACKGROUND:** Fibrous capsules (Fb) in response to cardiovascular implantable electronic devices (CIEDs), including a pacemaker (P) system, can produce patient discomfort and difficulties in revision surgery due partially to their increased compressive strength, previously linked to elevated tissue fibers.

**OBJECTIVE:** To quantify structural proteins, determine if biologic extracellular matrix-enveloped CIEDs (PECM) caused differential Fb properties, and to implement a realistic mechanical model.

**METHODS:** Retrieved Fb (-P and -PECM) from minipigs were subjected to biomechanical (shear oscillation and uniaxial compression) and histological (collagen I and elastin) analyses.

**RESULTS:** Fb-PECM showed significant decreases compared to Fb-P in: low strain-loss modulus (390 vs. 541 Pa) across angular frequencies, high strain-compressive elastic modulus (1043 vs. 2042 kPa), and elastic fiber content (1.92 vs. 3.15 μg/mg tissue). Decreases in elastin were particularly noted closer to the implant’s surface (Fb-PECM = 71% vs. Fb-P = 143% relative to dermal elastin at mid-tangential sections) and verified with a solid mechanics hyperelasticity with direction-dependent fiber viscoelasticity compression simulation (r^2^ ≥ 98.9%).

**CONCLUSIONS:** The biologic envelope composed of decellularized porcine small intestine submucosa ECM for CIEDs promoted fibrous tissues with less elastic fibers. Novel compression modeling analyses directly correlated this singular reduction to more desirable subcutaneous tissue mechanics.

## 1. Introduction

The formation of fibrous capsules is a foreign body reaction (FBR) against nondegradable materials, long-term implanted into the host system, dependent on factors including the material’s physical and chemical surface properties [1–4]. This tough tissue is induced to wall-off and systemically-isolate foreign objects (like cardiovascular implantable electronic devices (CIEDs) [5] with titanium housing and their polymer-coated leads) and is a general indicator for implant biocompatibility: the thinner the layer, the more compatible the material [3, 6, 7]. The fibrous capsule (also called fibrotic scar tissue or fibrosis) is distinct from other tissue types in the subcutaneous space and is mainly composed of compact collagen fibers produced by activated fibroblasts. The capsule can also contain layers of loose granulation tissue, immune cells, vessels, and a variety of enzymes and signaling molecules [1, 8, 9]. Aside from collagen (mainly type I, with occasional types II and III) [10–12], fibrous capsules have other extracellular matrix (ECM) structures, most notably elastic fibers [1, 9, 13, 14]. Elastic fibers are responsible for elastic recoil after stretching and compression for reversible recovery after deformation caused by applied loads, particularly beneficial for skin and blood vessels [15–17]. However, in fibrous capsules surrounding medical implants, their presence in excess may be detrimental to the tissue mechanics. An elastic fiber’s major component (~ 90%) is elastin. Initially, the relatively small (M_w_ ~ 60 kDa) soluble precursor tropoelastin is secreted by cells (such as fibroblasts), which then assembles and is crosslinked (forming covalent bonds) into elastic fibers via elastogenesis. Lysyl oxidase (and related enzymes) oxidize lysine residues for crosslinking with other tropoelastins and ECM proteins including collagens.

The fibrous capsule FBR is dynamic (time-dependent) and constantly remodeled depending on the stimuli. CIED-induced fibrous tissues can be used to an advantage: to subcutaneously affix and prevent device migration [18–21]. Conversely, too much fibrosis causes unwanted scar appearance, local tissue hardness, high compressive strength, and difficulties during CIED generator changes and pacing lead revision surgeries [22–26]. In patients with silicone breast implants, contracture, which induces increased stiffness, discomfort, and pain, can be attributed in part to increased elastin expression [13, 16, 27]. Similarly, issues encountered with CIEDs may be due to the overabundance of elastic fibers. Biologic ECM envelope products composed of decellularized porcine small intestine submucosa (SIS) were developed [28–30] to potentially mitigate the fibrous response. Besides acting as a cover for the CIED surface, SIS ECM implants are rich in bioactive growth factors, glycosaminoglycans, and structural proteins that can affect the subcutaneous microenvironment and FBR [31–35]. To investigate their underlying effect, this study compared fibrous capsule biomechanical and histological properties and associated elastic fiber distribution from pacemakers without and with biologic ECM envelopes in a short-term minipig subcutaneous model. A computer model was proposed and fitted to explain and verify the observed compressive stress-strain responses and correlate them to elastin amounts and arrangements.

## 2. Materials and methods

### 2.1 Animals and fibrous dissection

Surgical procedures and animal care were conducted at an independent research organization (American Preclinical Services, Minneapolis, MN). Protocols were approved by the Institutional Animal Care and Use Committee (IACUC), and animals received humane care in compliance with NIH guidelines [36]. Human pacemakers (pulse generators) with an attached, coiled pacing lead (Edora 8 SR-T and Solia JT 45, Biotronik, Lake Oswego, OR), without (P) and with (PECM) decellularized SIS ECM envelopes (CanGaroo^®^ Envelope (medium), Aziyo Biologics, Silver Spring, MD) were implanted subcutaneously (at 8 per group) in contralateral ventral neck-chest areas of adult miniature pigs (minipigs), then sacrificed and necropsied for a total of 3 months duration (based on [19]). Implants that failed before the endpoint were excluded from this study. After pacemaker retrieval (Fig. 1), the surrounding fibrous capsule (Fb) and adjacent adipose (Ad) with occasional underlying muscle tissue were excised, trimmed, and assigned as groups (Fig. 2): **Fb-P** (fibrous tissue from pacemaker only, n = 5) and **Fb-PECM** (fibrous tissue from pacemaker in ECM envelopes, n = 2), with Ad (adipose, n = 4) control, where n = biological replicates (individual animals). Multiple technical replicates (≥ 3) were obtained per biological replicate per test to account for individual variability [37].

**Fig. 1.**
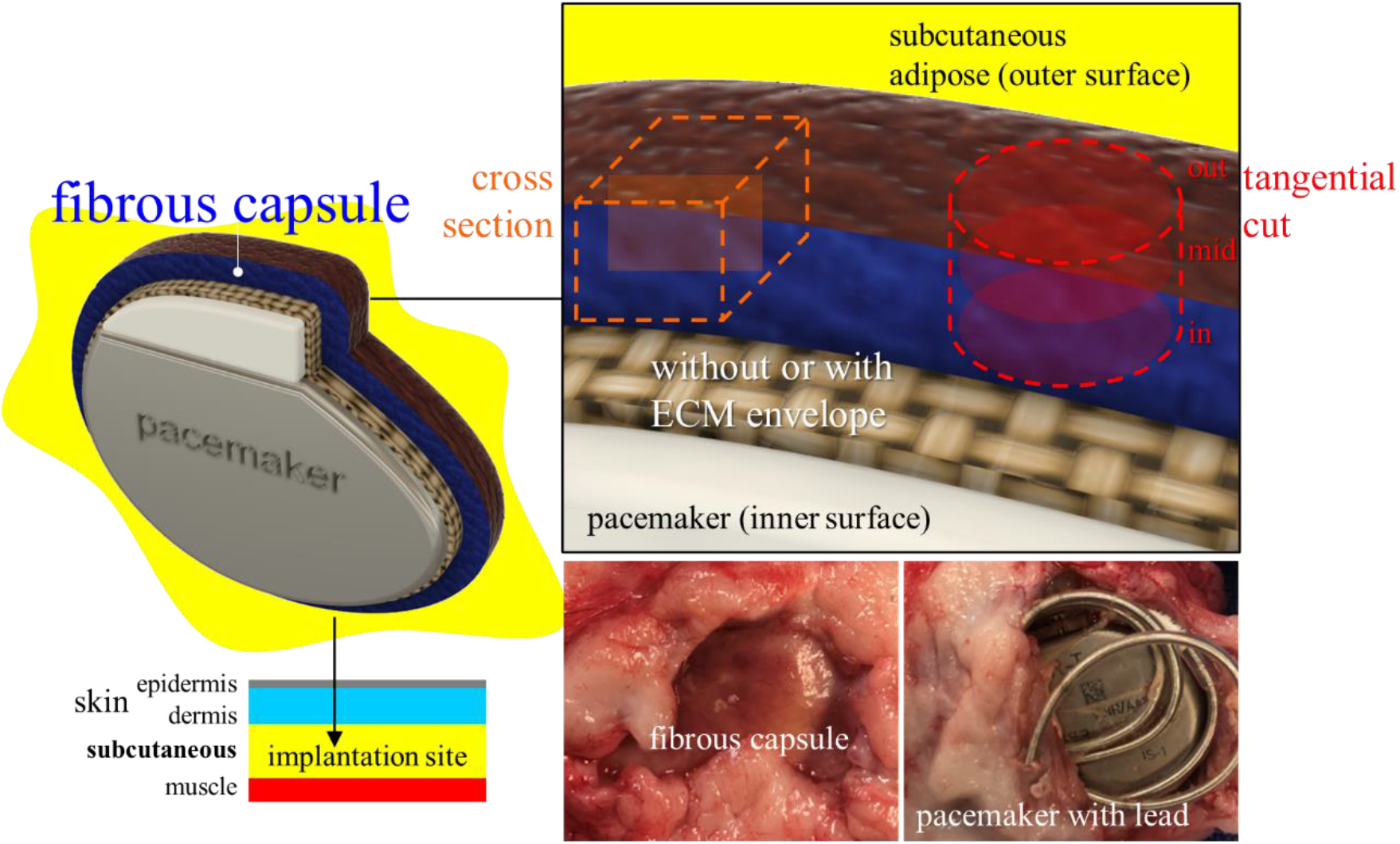
A pacemaker model (top left) implanted in the subcutaneous pocket (bottom left) with the subsequent fibrous capsule (Fb) response (bottom center). After retrieval (bottom right), select Fb with distinct inner and outer surfaces was cut into cross and tangential (in, mid, and out) sections (top right).

**Fig. 2.**
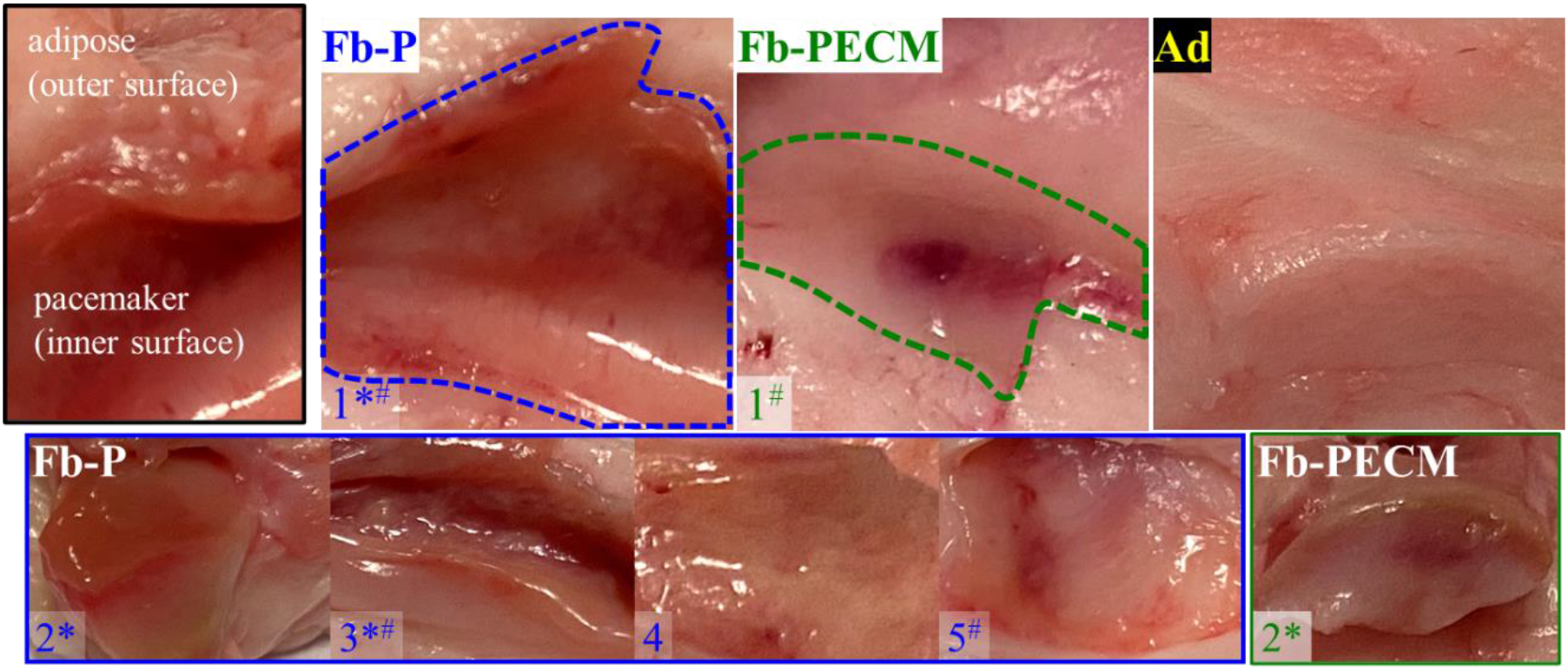
Images of Fb-P (fibrous tissue from pacemaker only, n = 5), Fb-PECM (fibrous tissue from pacemaker in ECM envelope, n = 2), and Ad (adipose control) surrounding tissues. Fibrous tissues are located inside the dashed outlines for both samples labeled “1”. *Fb-P samples: 1-3 and Fb-PECM: 2 had pronounced fibrous connective tissue appearance. #Fb-P: 1, 3, and 5, and Fb-PECM: 1 had some regions of hematoma.

### 2.2 Shear oscillation rheology

Tissues were subjected to rotating shear oscillation at exponentially increasing angular frequencies (ω) per fixed maximum shear strain (γ) to obtain their viscoelastic properties: storage modulus (G’), loss modulus (G”), delta or shear stress to strain phase shift (δ) = tan^-1^(tan(δ)) in degrees from 0° (pure solids) to 90° (pure liquids), and dynamic modulus (magnitude of complex modulus (G* = G’ + iG”) = |G*|) = (G’^2^ + G”^2^)^0.5^ = G’/cos(δ). Cylindrical biopsies at 8 mm-diameter (d) (Fb inner surface down and outer surface up) were tested in DHR-2 (Discovery Hybrid 2, TA Instruments, New Castle, DE) using parallel plates with a probe d = 12 mm and gap assigned to 2 mm. Oscillation experiments were performed at 5% (low) and 40% (high) γ from 1-40 rad/s (0.16-6.34 Hz) ω.

### 2.3 Axial compression

Unconfined uniaxial compression was conducted to obtain [38]: true compressive stress (σ), strain (ε), and elastic modulus (E) = Δσ/Δε at 0-10% (low) and high 30-40% (high) ε. Samples with d = 8 mm and height (h) = entire Fb thickness were placed onto the baseplate (Fb inner surface down and outer surface up) and compressed with a 6 mm chamfered indenter at 0.1 mm/s until ~ 40-60% ε (Instron 3345, Norwood, MA). Instantaneous E = dσ/dε = numerical derivative of σ with respect to ε was reported (centered finite difference with a 2-term Taylor: (dσ/dε)_i_ = (-σ_i+2_ + 8σ_i+1_ – 8σ_i-1_ + σ_i-2_)/12(ε_i_ – ε_i-1_) [39]) to better display changes in σ = *f*(ε).

### 2.4 Soluble elastin

Fastin^™^ elastin assay (Biocolor, Carrickfergus, Northern Ireland, UK) [40, 41] quantified elastic fibers (EFs) via released water-soluble α-elastin from tissues. Fresh specimens were processed through α-elastin extraction (100 °C in 0.25 M oxalic acid), precipitation, and complexation with 5,10,15,20-tetraphenyl-21H,23H-porphine tetrasulfonate (TPPS). The complex was recovered, TPPS released, absorbance at 513-nm measured, converted to mass using standards, and normalized to tissue mass (compared to a pig skin positive control). The assay was deemed reliable if the obtained ratio of elastin in the skin control was around the expected 3.5 μg/mg [40].

### 2.5 Tissue staining

Fb samples were processed for histology to determine microscopic thickness, morphology, and structure using Masson’s trichrome [42] (MT, Trichrome Stain Kit ab150686, Abcam, Waltham, MA) [43] for connective tissue components (cell nuclei = purple, cytoskeleton = red, and tissue collagen = blue), Van Gieson’s elastic (VGE, Elastic Stain Kit ab150667, Abcam) [44] for EFs (black strands), and immunohistochemistry (IHC) for detection (brown signals) of collagen type I (1:100 anti-human collagen I alpha-1 1200-1450 fragment (ab233080, Abcam) compatible to pig elastin [45]) and EF elastin (1:200 anti-pig and human elastin (ab21610, Abcam)) [41, 46, 47]. The 3D differential distribution of elastin was investigated using four 2D: one cross and three tangential (inner (in), middle (mid), and outer (out) surface) sections (Fig. 1).

Briefly, tissues were fixed in 10% neutral buffered formalin, stored in 70% ethanol (EtOH), trimmed, dehydrated in increasing EtOH then xylene, and paraffin-embedded (Paraplast X-TRA^®^, Sigma-Aldrich, St. Louis, MO). Paraffin blocks were sectioned (at 5-μm), placed on slides, and rehydrated. After staining with MT and VGE, samples were dehydrated and soaked in Histomount (Thermo Fisher Scientific, Waltham, MA). For IHC, rehydrated sections (1 negative (−) control and ≥ 1 experimental/slide) with dried boundaries were circled with a hydrophobic barrier (ImmunoPen, Sigma-Aldrich). Endogenous peroxidase inhibition and antigen retrieval (citrate pH 6 at 95 °C for 10 min) was performed before blocking with 10% normal goat serum. Specimens were hybridized with 1° antibody (Ab) for 2 hours (while blocking buffer alone was used for (−)), processed with rabbit specific HRP/DAB (ABC) Detection IHC Kit (ab64261, Abcam) [48], and mounted in ImmunoHistoMount (Sigma-Aldrich). Human skin (Sigma-Aldrich and Aziyo Biologics) (comparable to pig skin [49]) and ECM envelope (Aziyo Biologics) samples were included as controls.

Images were captured in Cytation 5 (Agilent, Santa Clara, CA) and processed with ImageJ (NIH, Bethesda, MD). IHC area ratios were quantified in thresholded and leveled images using ImageJ’s analyze particle area sum over polygonal tissue area selection. Individual slide (−) signals were subtracted.

### 2.6 Modeling of compression

COMSOL Multiphysics (COMSOL, Stockholm, Sweden) using the solid mechanics interface with nonlinear structural materials module was employed to generate a 3D time-dependent compression model matching the experimental using parameters: Fb (E = 295 kPa (determined through parametric sweep), density (ρ) = 1.05 g/mL, and Poisson’s ratio (ν) = 0.4999) and EF properties (E = 400 kPa (calculated based on [50]) and ρ = 1.2 g/mL). Fb was assumed as neo-Hookean hyperelastic incompressible [51]. EFs were modeled with direction-dependent (tangential and axial) domain node fibers (linear elastic with a volume fraction or ratio (r_v_)). The tissue was divided into sections (in, mid, and out) and each assigned with fibers with viscoelasticity of a single-branch generalized Maxwell containing: energy factor (f_e_) and relaxation time (t_r_) [50]. A rigid indenter at 0.1 mm/s prescribed velocity compressed Fb with constrained bottom boundary to 40%. Unstructured uniformly-distributed quadrilateral meshes along h were used. Average of the domain’s top boundary in contact with the probe was evaluated for stress = Gauss point von Mises stress and strain = displacement/h.

### 2.7 Data analysis

Data were processed in Excel (Microsoft) and MATLAB (Mathworks, Natick, MA). Technical replicate values were averaged, and biological replicates’ values again averaged then reported as mean ± standard deviation. Student’s t-test and analysis of variance (ANOVA, 1 and 2-factor) with Tukey-Kramer were used for comparison using a probability (p) < 0.05 deemed significantly different and represented as: ***p < 0.001, **p < 0.01, and *p < 0.05.

## 3. Results

### 3.1 Fibrous capsule gross evaluation

Implants induced Fb tissues (Fig. 1) in the subcutaneous region. The fibrous appearance was prominent in 3/5 (60%) and 1/2 (50%) for Fb-P (samples 1-3) and Fb-PECM (2), respectively (Fig. 2). These tissues varied in thicknesses in situ from ~ 2-6 mm within each sample and across biological replicates. They were attached to adipose (Ad) tissues on their outer surfaces, while their inner surfaces (adjacent to implant) were smooth, slippery, and lined with interstitial fluid. Localized blood patches (subcutaneous hematoma) were observed on inner sides in 3/5 (60%) and 1/2 (50%) of Fb-P (samples 1, 3, and 5) and Fb-PECM (sample 1), likely due to the surgical procedure. No signs of swelling were noticed.

### 3.2 Response to periodic shear

Fb tissues subjected to 5% γ oscillation showed G’ > G” (Fig. 3a), indicating a more solid elasticity with δ < 45° (Fig. 3c), generally independent of ω. G’ and G” displayed parallel plots due to their undamaged structures (nondestructive condition). The deformation was small to remain in the linear viscoelastic region (where data deemed reliable). The |G*| (resultant of G’ and G”) for Fb (P vs. PECM) were similar (p = 0.063), but fibrous capsules had lower (***p < 0.001) |G*| than adipose tissues (Fig. 3b); showing these connective tissue types can easily be distinguished. Importantly at **low γ, Fb-PECM’s G” was statistically lower (*p = 0.029) compared to Fb-P’s** (Fig. 3a, Table 1).

**Fig. 3.**
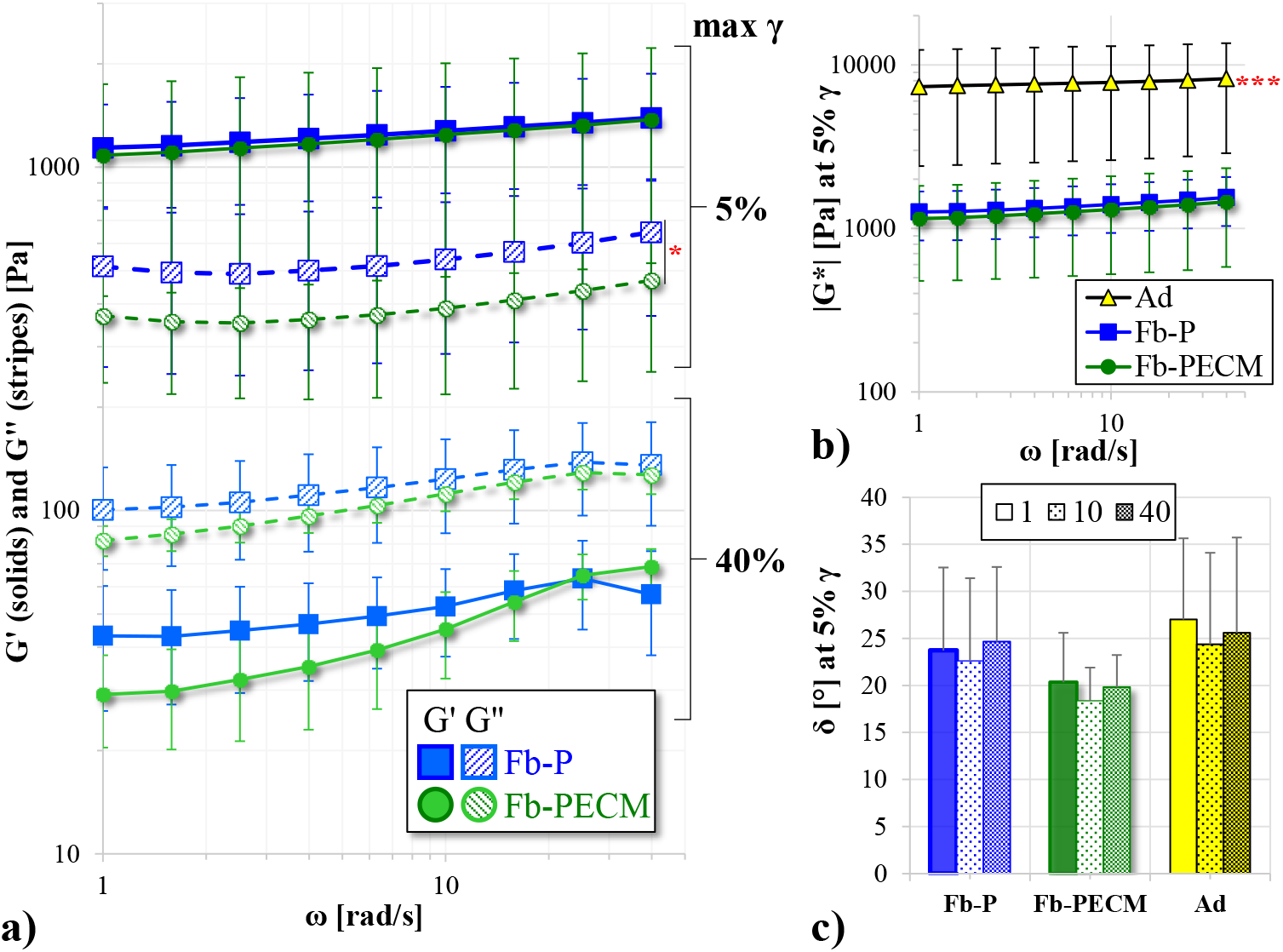
a) Storage (G’) and loss (G”) moduli vs. angular frequency (ω) in a log-log plot at 5% and 40% maximum shear strain (γ) of Fb-P (blue) and Fb-PECM (green). G”, 5% γ: Fb-P > Fb-PECM (*p < 0.05). b) Log-log (5% γ) of dynamic modulus (|G*|) vs. ω. Ad (yellow) > Fb (***p < 0.001) c) Phase shift (δ) bar at 5% γ and 1, 10, and 40 rad/s ω.

**Table 1.**
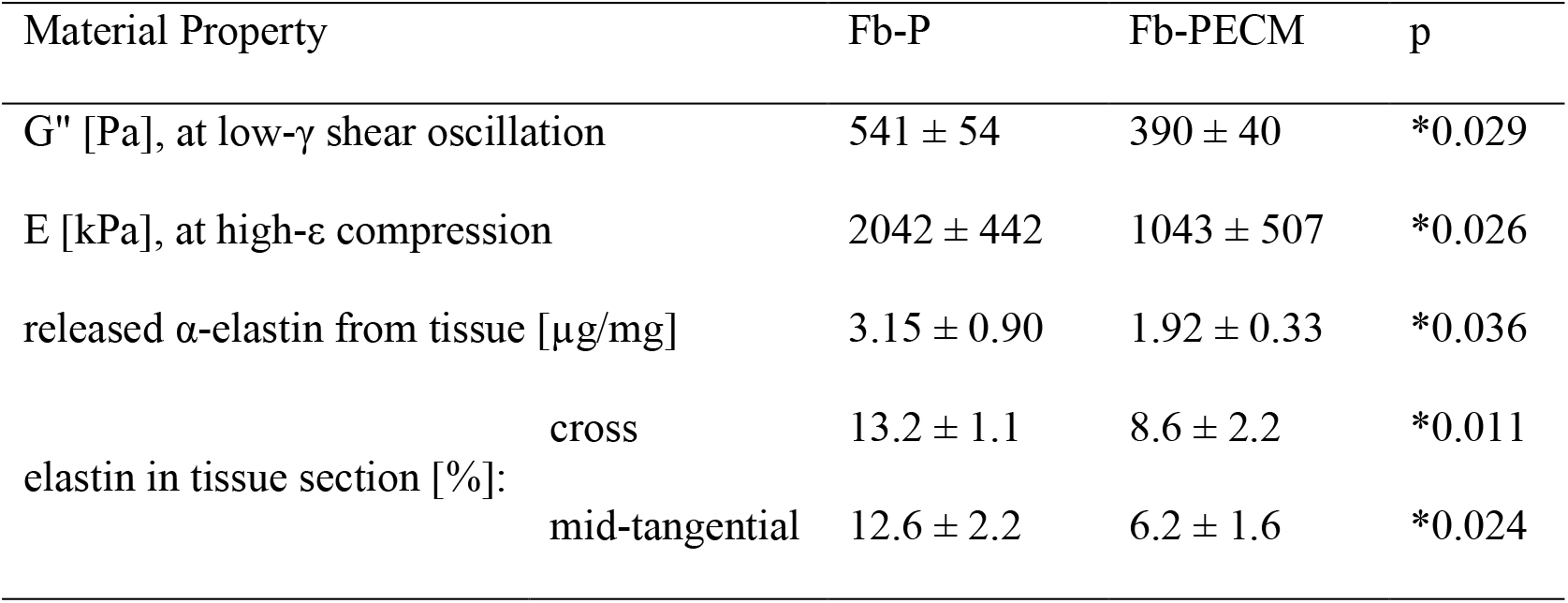
Fibrous (Fb) tissue properties with significant difference.

At 40% γ, G’ trended at < G” (Fig. 3a) and δ > 45° (49-67° spread), showing materials with dominant fluid-like and viscous properties. The high γ led to stretching which appeared to release bound water molecules and influence the elevated loss modulus. Pairwise G’ and G” demonstrated similarities despite an apparent lower pattern for Fb-PECM.

### 3.3 Effect of compression

Macroscopic thicknesses (for strain calculation) using a caliper yielded 4.0 ± 1.1 for Fb-P and 3.3 ± 0.5 mm for Fb-PECM, but still statistically similar (p = 0.59).

Compression to high strains (some up to ~ 60%) led to reversible deformation within the elastic region even with nonlinearity of the σ-ε curves. Ad had drastically lower σ and its derivative (dσ/dε) compared to both Fb’s (Fig. 4a-b), which rendered to lower (***p < 0.001 at 0-10% and at least *p < 0.05 at 30-40% ε) compressive E (Fig. 4c).

**Fig. 4.**
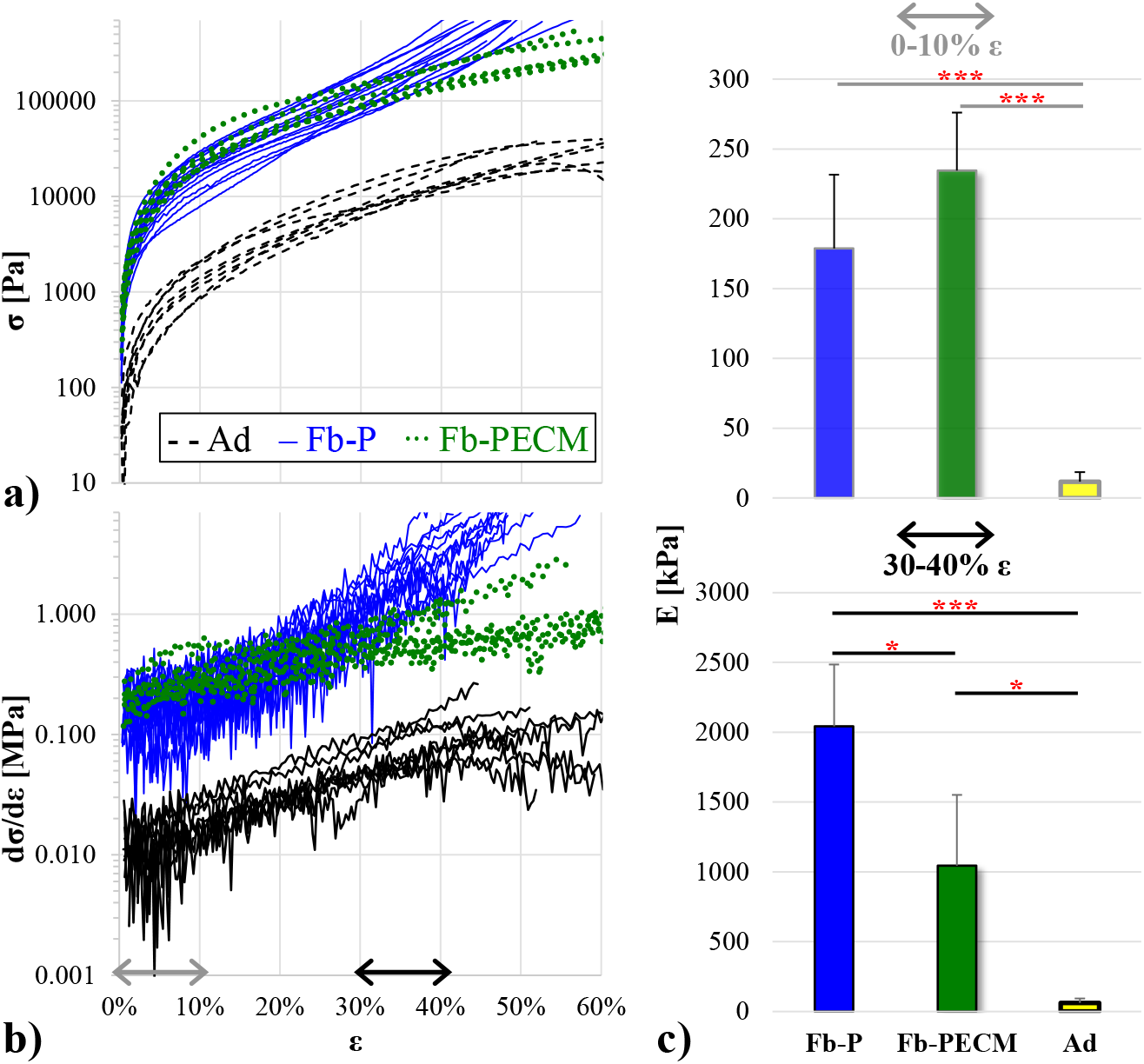
a) True compressive stress (σ) vs. strain (ε) in semi-log of Fb-P (blue), Fb-PECM (green), and Ad (black/yellow) technical replicates. Ad trended < Fb. Fb-P’s curvature looks different compared to Fb-PECM and Ad’s, highlighted in the next graph. b) Semi-log of instantaneous elastic modulus (dσ/dε) vs. ε, indicating 0-10% (gray) and 30-40% (black double arrows) regions. c) Compressive elastic modulus (E) at 0-10% (top) and 30-40% (bottom) ε. Pairwise significance: *p < 0.05 and ***p < 0.001, notably: Fb-P > Fb-PECM at 30-40% ε.

It is remarkable that Ad and Fb-PECM’s curves had generally similar semi-log shapes (with adipose just translated downwards), while Fb-P’s exhibited a distinct pattern (Fig. 4a) and more noticeable in numerical differentiation (Fig. 4b blue curves). Quantitatively **at high ε (30-40% or higher), Fb-P had significantly greater (*p = 0.026) slope or stiffer E than Fb-PECM** (Fig. 4c, Table 1).

### 3.4 Amount of α-elastin

Fibrous tissues were detected to contain more (**p = 0.002 for Fb-P and *p = 0.011 for Fb-PECM) elastin compared to adipose (Fig. 5). Importantly, **Fb-P had statistically higher (*p = 0.036) tissue elastin than Fb-PECM** (Fig. 5b, Table 1), indicating observed biomechanics differences were influenced by elastic fibers.

**Fig. 5.**
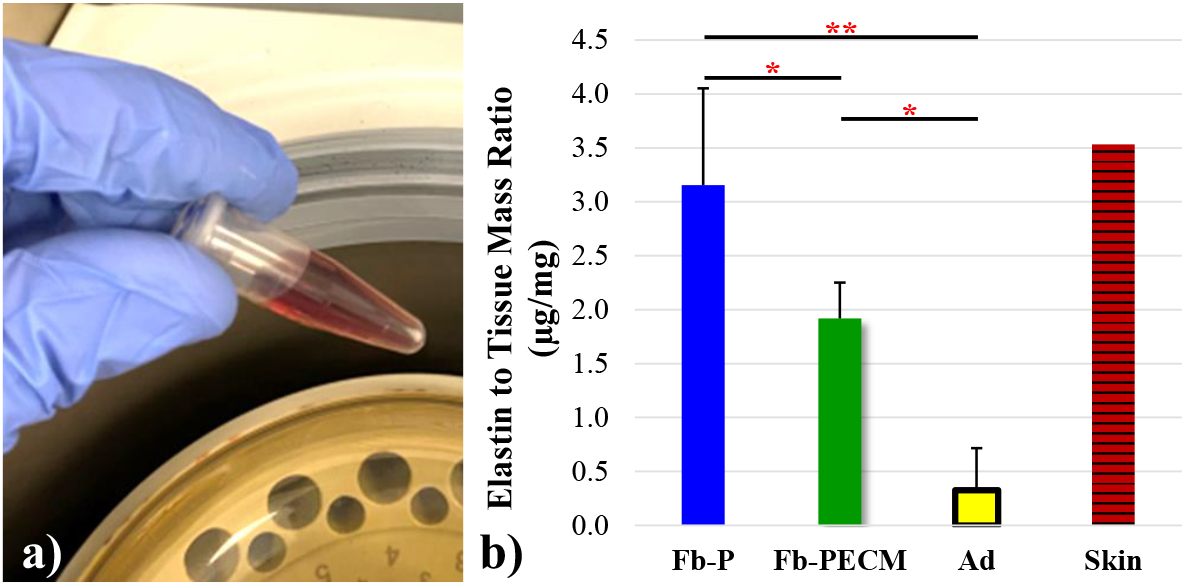
a) Reddish-brown TPPS after release from α-elastin pellet. b) Normalized elastin-to-tissue mass ratio of the two Fb groups, Ad, and a skin control. Pairwise significance: *p < 0.05 and **p < 0.01, importantly: Fb-P > Fb-PECM.

### 3.5 Histology analysis

Sections of the samples from both Fb groups showed collagen secreted by activated fibroblasts, filling the entire thickness with interspersed blood vessels (Fig. 6). Fb-PECM stained their cell nuclei faint and ECM lighter even after several repeats, perhaps due to differences in chemistry. Additionally for Fb-PECM, there were no signs of remnant ECM envelope suggesting full remodeling of the SIS. No foreign body giant cell was noticed indicating the absence of chronic inflammation. In some sections, adipose tissue can be seen in the mid to outer zones, which indirectly indicated fibrous formation originating from the implant’s surface. Regions with hematoma demonstrated reddish coloration in MT with round nuclei and brownish in VGE, likely as macrophages and blood, found in the inner surface.

**Fig. 6.**
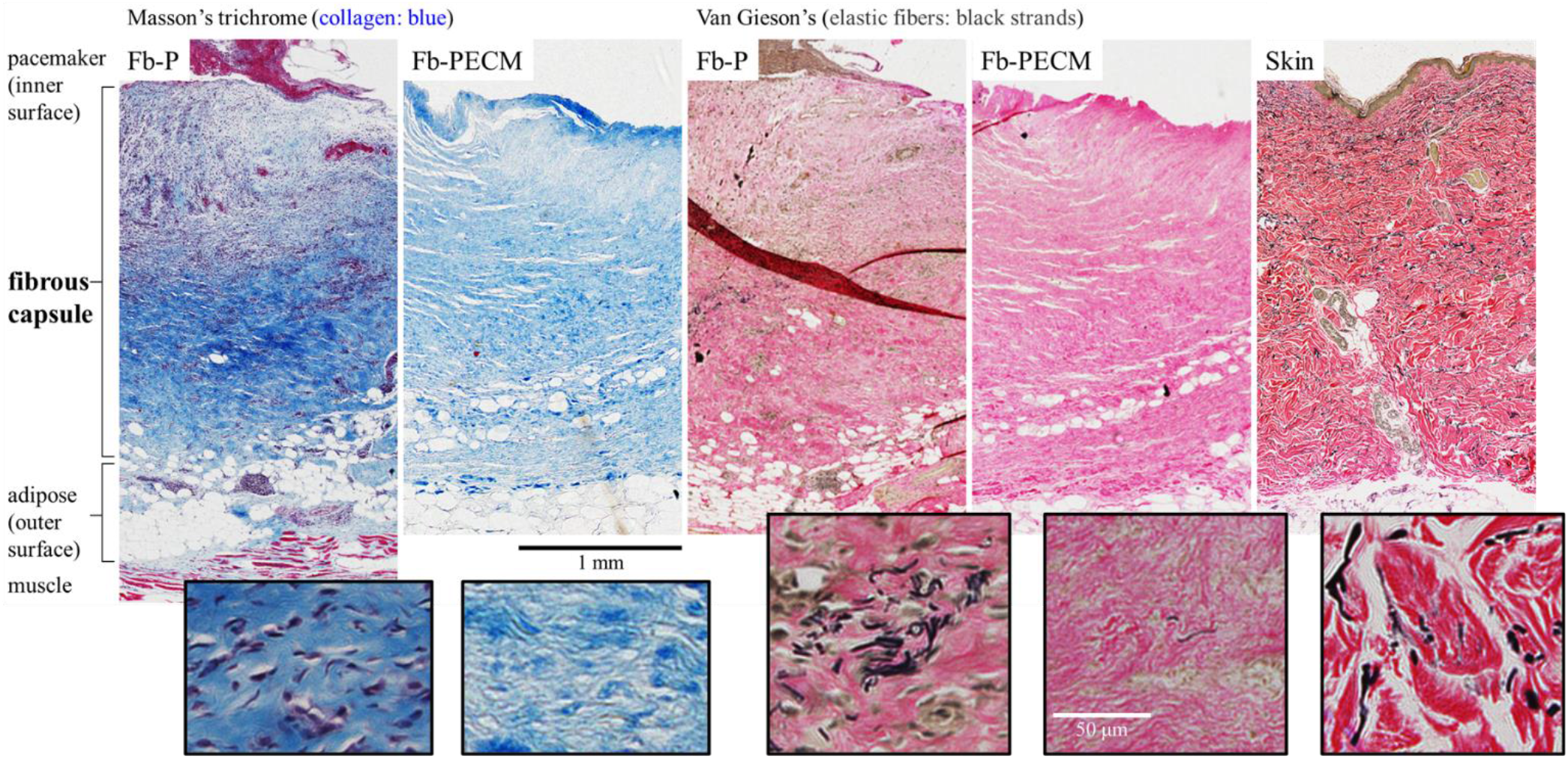
Connective tissue stains: Masson’s trichrome (MT) and Van Gieson’s (VGE) of representative Fb cross-sections with layers spanning from the implant-adjacent inner to the outer surface attached to Ad. Human skin VGE with epidermis and dermis tissues included as controls. Boxed insets demonstrate magnified regions with fibroblasts, collagen ECM (blue in MT), and elastic fibers (black strands in VGE).

Microscopic thicknesses (2.7 ± 1.1 for Fb-P, 2.2 ± 0.3 for Fb-PECM, and 2.3 ± 0.1 mm for skin) were statistically similar (p = 0.89). Interestingly, the overall morphology of fibrous capsules resembled the skin dermis (Fig. 6 VGE).

Elastic fibers (EFs) were detected as black strands and visually more present in cross-sections of Fb-P, especially in regions closer to the inner surface (Fig. 6 VGE inset). Immunostaining confirmed **higher (*p = 0.011) differentially-expressed EF elastin in Fb-P vs. Fb-PECM** (Fig. 7, Table 1, 1.49 ± 0.12 and 0.98 ± 0.25 relative to dermis control levels). In tangential cuts (Fig. 7a boxed), there appeared to be more elastin in Fb-P (and in dermis) vs. Fb-PECM at inner and mid zones. Statistically, **Fb-P > Fb-PECM (*p = 0.024) at mid-tangential** (Fig. 7b, Table 1). Fb sections showed various stages of elastogenesis, ranging from diffuse elastin staining to assembled EFs with different orientations. The biologic ECM envelope displayed lower (*at least p < 0.05) background elastin compared to others.

**Fig. 7.**
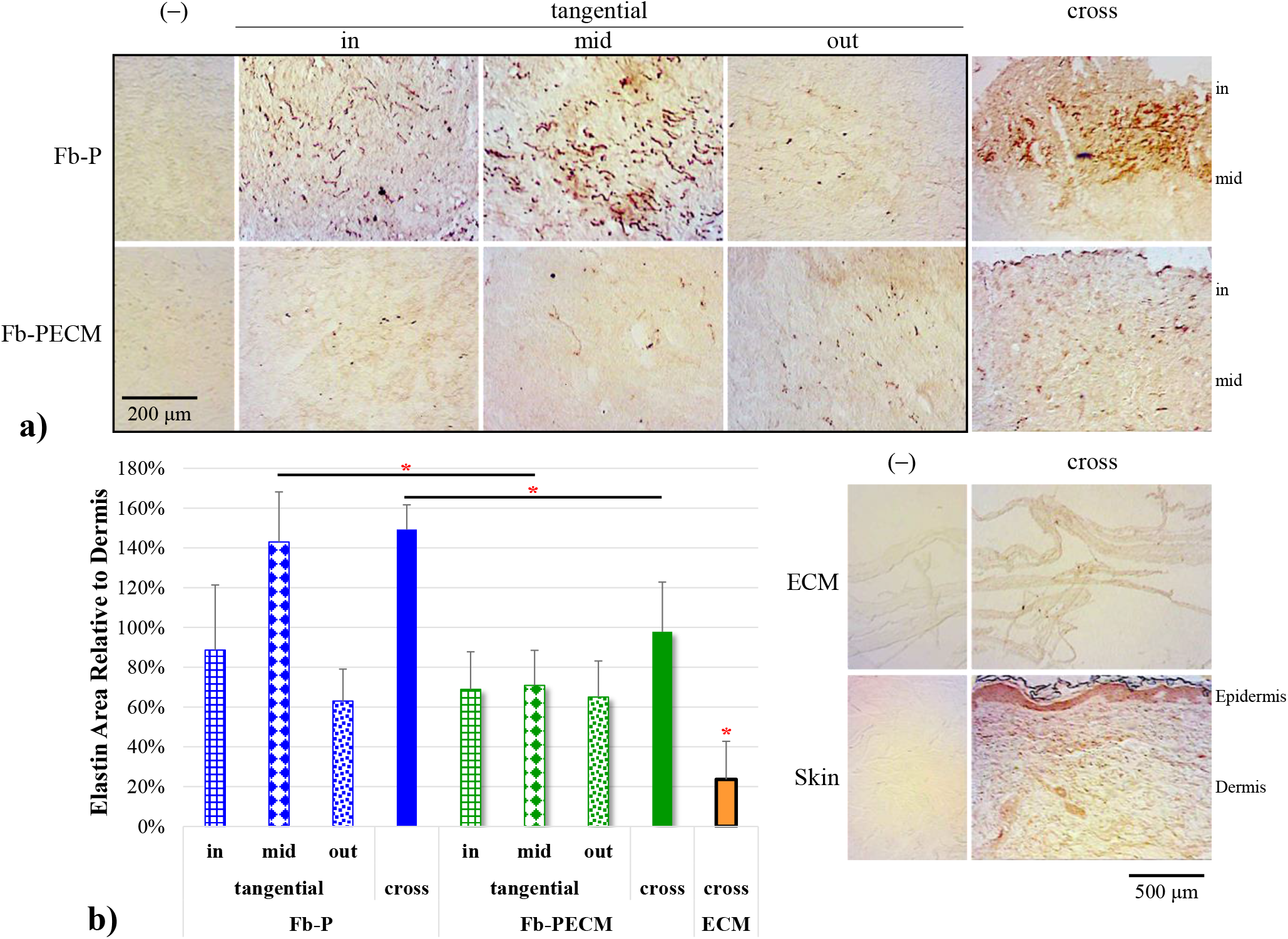
a) Representative elastin-IHC sections with negatives (–) from the two Fb groups and biologic ECM envelope and skin (showing epidermis and dermis tissues) controls. Visually, Fb-P > Fb-PECM for in- and mid-tangential and cross-sections staining. b) Elastin in tissue area ratio compared to dermis bar graph. Mid-tangential and cross-sections: Fb-P (blue) > Fb-PECM (green) (*p < 0.05). ECM envelope (orange) < all other groups (*p < 0.05).

Collagen type I is prominent among all tissue groups: Fb-P, Fb-PECM, ECM envelope, and dermis controls were found to be statistically similar (p = 0.81) (Fig. 8). Its density was qualitatively more intense in bundles observed in the dermis (Fig. 6, Fig. 8a). Conversely, collagen I in fibrous capsules did not appear bundled but as separate fibers, suggesting less organization and maturation. EFs did not colocalize with collagen I in Fb samples.

**Fig. 8.**
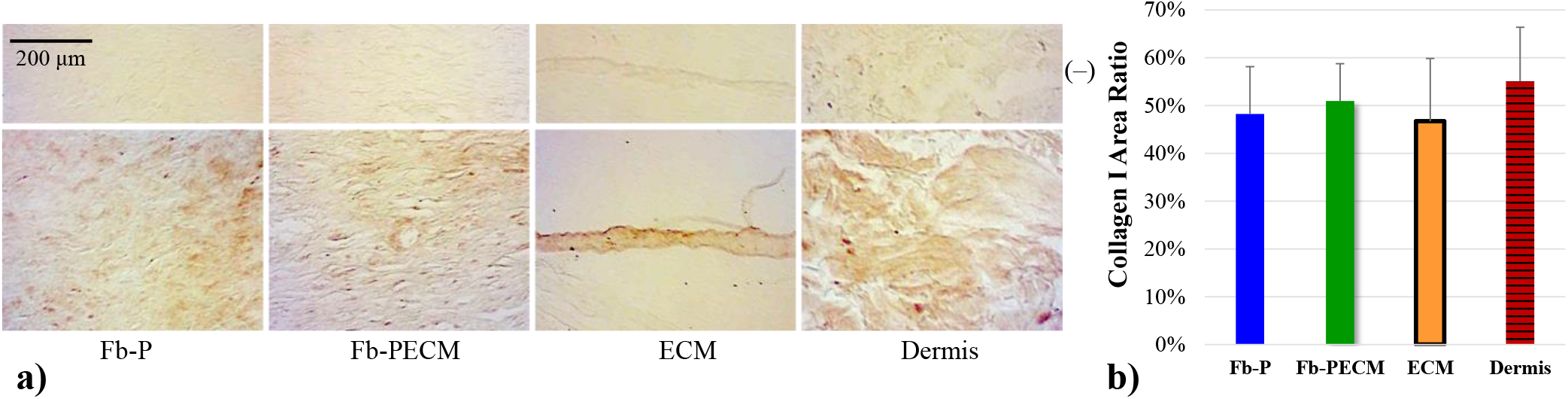
a) Collagen I-IHC cross-sections from Fb groups, ECM envelope, and skin dermis. b) Bar of collagen I in tissue area ratio with 47-55% range with no significant differences between groups.

### 3.6 Computer simulation

Simulation with 6-fold axially-oriented fibers at the inner and mid-Fb (in + 1/3 mid) relative to 12% rv of tangential whole-tissue fibers (with consistent EF properties) resulted in a compressive σ-ε curve that agreed well (r^2^ = 99.7%) compared to experimental Fb-P observation, including elevated von Mises stress seen across tissue equivalent to higher compressive strength (Fig. 9a). When these axial elastic fibers were removed (at 0 fiber, Fig. 9b-c), the plot fitted accurately to Fb-PECM experimental (Fig. 9d, r^2^ = 98.9%). Different iterations of domain material models were simulated (Fig. 9e), but ultimately the hyperelastic with regional viscoelastic direction-dependent fibers showed the best for modeling Fb-P and Fb-PECM which agreed with experimental trends relating compression (Fig. 4a, Fig. 9d) to differential elastic fiber expression (Fig. 5b, Fig. 6, Fig. 7) using the viscoelasticity parameters: f_e_ = 2% and t_r_ = 50 s.

**Fig. 9.**
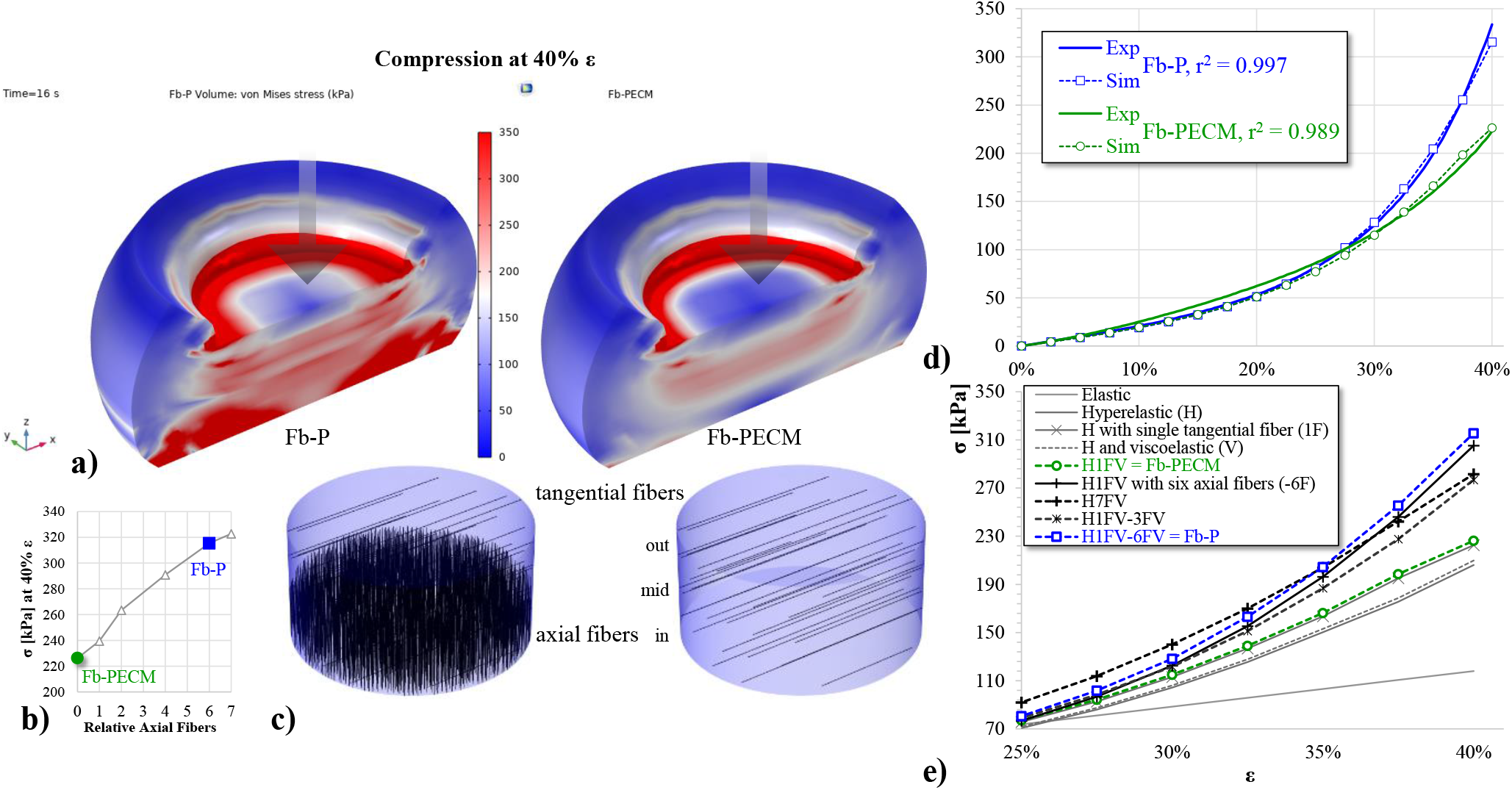
a) Graphical 3D of von Mises stress and deformation of Fb compression simulation at 40% ε. b) Compressive stress (σ) vs. axial fiber amount, indicating 0 for Fb-PECM and 6 for Fb-P. c) Locations and directions of fibers in Fb domains. d) Nonlinear scatterplot of σ vs. ε, displaying good agreement between Fb experimental (Exp) and simulation (Sim) values. e) σ vs. ε Sim curves of various material models.

## 4. Discussion

### 4.1 Biologic ECM envelopes mitigate elastic fiber formation and fibrous tissue stiffness

The fibrous capsule response in this study was influenced by host contact to implant surface materials: titanium housing of pulse generator, epoxy resin of connector receptacle, silicone rubber of sealing plug, silicone rubber and polyurethane cover of pacing lead, and decellularized porcine SIS ECM of the envelope [28, 52]. Macro and micro-histologic features of recovered and processed fibrous tissues, without or with the biologic ECM envelope treatment, indicate differing degrees of FBR biocompatibility, previously observed in a rabbit model [53]. As expected from prior animal studies, both implant groups induced varying levels of fibrous tissues prominent in collagen I [10–12, 54]. Despite similarities in appearance, thickness, and collagen type I morphology and content, we found five properties with statistically less magnitudes from Fb-PECM samples, summarized in Table 1.

Changes in biomechanics at specific strain conditions, as seen at low-γ shear oscillation and high-ε compression, can be explained by the anisotropic [55] levels of tissue elastin, which constitute the mature functional form of load-bearing elastic fibers. Elastic fibers are abundant in the capsular tissue from several other different subcutaneous implant studies [1, 9, 13, 14]). High-strain E of Fb-P tissue samples compressed at ~ 30% were well within the 1.6-5.7 MPa of silicone implant fibrous capsules collected from breast capsular contracture patients [13]. E values for Ad control tissue fell within the acquired spread from human adipose [56], supporting that our tests were sound and accurate. Use of the biologic ECM envelope significantly lowered Fb-ECM levels of elastin and elastic fibers versus Fb-P, particularly in tissue regions close to the pacemaker and leads. The reduction strongly correlated with a drop in E below Fb-P and the previously reported range of undesirable tissue from breast capsular contracture. Since this study only had a short 3-month duration, mechanical differences between the Fb-ECM and Fb-P groups should widen over time as elastic fibers continue to accumulate at a faster rate in the Fb-P group, pushing the mechanical properties to match tissue from the most severe cases of contracture. These data suggest biologic ECM envelopes could be used to alter the FBR for subcutaneous implants and mitigate excess accumulation of structural components, like elastic fibers, in the fibrous tissue that increase tissue elastic modulus and cause contraction.

Reducing tissue stiffness following subcutaneous implantation surgeries may improve patient comfort and long-term clinical outcomes. For patients receiving CIEDs, a hard fibrotic tissue can cause lead dislodgment [57, 58]. Tissue with higher stiffness values encasing generator and/or leads also increases the difficulty of revision procedures and contributes to higher complication rates observed for revision procedures compared to those reported for de novo implantations [24, 25, 59, 60]. Using biologic ECM envelopes to wrap CIEDs and modify the FBR could be an effective method to reduce these complications and improve patient health.

### 4.2 Novel compression modeling effectively simulates fibrous tissue from CIED FBRs

In compression testing of fibrous capsules reported in previous studies, excess collagen levels confounded the mechanical effects of elastic fibers. With statistically similar levels and orientation of collagen between this study’s two tissue groups, the role of elastic fibers could be isolated in computer modeling. In the proposed hyperelastic model with viscoelastic fibers (Fig. 9), Fb-PECM was set as the starting fibrous tissue baseline, assuming at the early timeline of Fb-P’s development that ultimately produced more elastic fibers. For this domain baseline, the simulated compressive stress-strain curve agreed with experimental once elastic fiber orientation was assigned as force-orthogonal (tangential or radial) and its tissue volume fraction to 12%, within acceptable degree of variability to the observed 8.6% (in cross-section). Addition and successive increase of force-parallel (axial) fibers to 6 times greater in the inner 44% region then created the accurate Fb-P compression model. The fiber ratio ultimately plateaued to around 7-fold maximum capacity, indicating peak resemblance to the dermal control and correlating with the similarities in elastin quantity and histology, which possibly signifies a similar tissue genesis. It is noted that fibers placed closer to the implant provided better simulation outcome with higher stress-strain influence than the mid and outer zones. Overall, evidence considering statistics, potential sources of error, visual, and modeling analyses suggest that more elastic fibers are found toward the CIED surface in Fb-P. This also resulted in higher loss modulus taken in the rheology experiment due to elastic fiber-dependent increased water absorption and tissue retention [16, 61].

## 5. Conclusions

CIEDs wrapped in the biologic ECM envelope reduced elastic fibers in the fibrous capsule region close to the implant, which directly led to a more subtle biomechanical response including lower compressive strength with some likeness to surrounding adipose tissue. Novel mechanical modeling highlighted the potential of this envelope for improved clinical outcomes of CIED implantations. A long-term study with more biological replicates for investigating elastic fiber anisotropy and associated cells and structures can be conducted in the future.

## Acknowledgements

We thank our research group undergraduates: Mariana Cabral, Megan Forst, Jayda Lewis, Andrew Tarabokija, Henna Chaudhry, and Evan Carroll for their lab assistance and feedback, Dr. Michael Dores (Biology) for help and access to imaging resources, and Engineering department and Hofstra University staff, especially Lori Castoria, Lynne Espiritu, Liz Downey, and Dean Sina Rabbany for overall support.

